# Role of Prefrontal Cortex during Sudoku task: fNIRS study

**DOI:** 10.1101/2020.05.25.115121

**Authors:** Patil Ashlesh, K K Deepak, Kochhar Kanwal Preet

**Author notes:** Corresponding Author: Prof. Kochhar Kanwal Preet, M.D., DNB, Ph.D, MNAMS, Cognitive Neurophysiology Lab, Department of Physiology, All India Institute of Medical Sciences, New Delhi, India -110029.

## Abstract

Sudoku is a popular leisure time activity that involves no math, but is based on logic based combinatorial number placement in a matrix. Many studies have been dedicated towards finding an algorithm to solve Sudoku but investigation of the neural substrates involved in Sudoku has been challenging. It is difficult to measure the brain activity during 9×9 Sudoku using traditional fMRI technique due to the procedural constraints. 16 optodes fNIRS (functional near infrared spectroscopy) forms an excellent alternative to study the activity of prefrontal cortex (PFC) during Sudoku task. Sudoku task was divided into two steps to understand the differential function of the PFC while applying heuristic rules. Classical two-way ANOVA as well as General Linear Model based approach was used to analyze the data. 28-noise free recording from right-handed participants revealed increased activity in all 16 optode locations during step 1 (3 × 3 subgrids) and step 2 (easy level 9×9 Sudoku) as compared to rest. Contrasting the step2-step1 revealed that medial regions of PFC were preferentially activated. These findings suggest the role of these regions, while applying multiple heuristic rules to solve 9×9 Sudoku puzzle.

**Graphical abstract:** 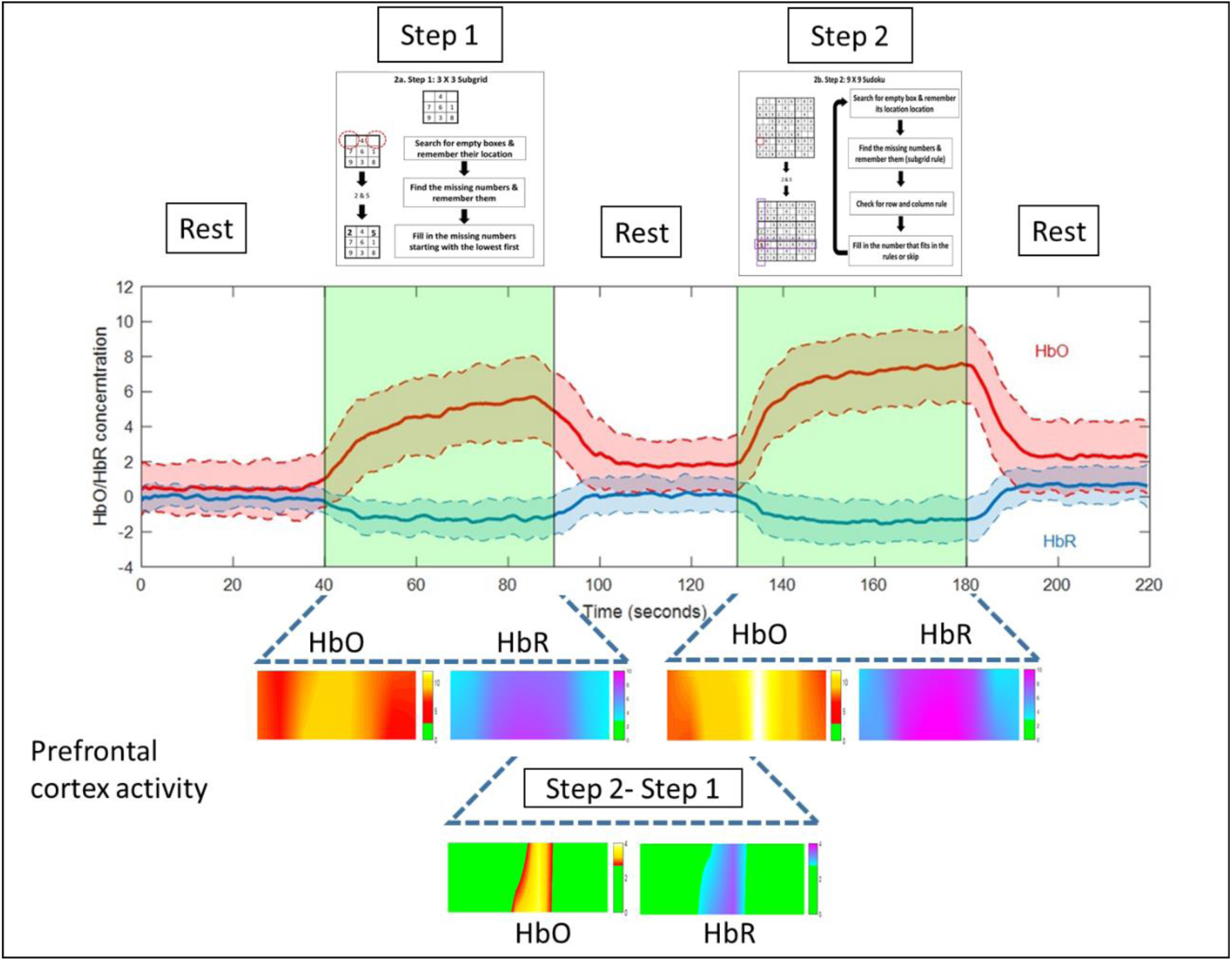

**Highlights:**

- This is first fNIRS study that tried to unravel the role of PFC during Sudoku task.
- Uniquely divided the Sudoku task into two steps to understand the differential role of PFC while applying multiple heuristic rules.
- Both the medial and lateral regions of PFC are activated during Sudoku task.
- However, the medial regions of PFC play a differential role, especially when we consider the row and the column rule of Sudoku.

## 1. Introduction

One of the commonest leisure time activities for all ages is solving puzzles. Popular amongst these puzzles is Sudoku, a logic based combinatorial number placement problem composed of a matrix with rows *(n*^*2*^), columns *(n*^*2*^), and subgrids *(n x n)* (Bargagliotti, 2006). The problem is based on three simple rules that the numbers should not repeat in the subgrid, row, or column. The commonest Sudoku puzzle is a 9 × 9 matrix with subgrids of 3 × 3. Sudoku is a good cognitively stimulating leisure time activity (Ferreira et al., 2015) and it even increases openness to new experiences affecting the personality trait of the individual (Jackson et al., 2012). Sudoku requires attention of the subject to analyze the grids and fill in the numbers, basically it requires no math but is based on logic (Bargagliotti, 2006). Many studies have been directed towards finding an algorithm to solve Sudoku, but investigation of the neural substrates involved in Sudoku has been challenging. Simplified version of Sudoku (4×4 matrix) has been used in various fMRI studies (Long et al., 2012; Qin et al., 2012; Wang et al., 2009; Xiang et al., 2009; Zhou et al., 2012). Qin *et al*. found involvement of left prefrontal cortex, dorsal anterior cingulate cortex and bilateral involvement of posterior parietal cortex, caudate nuclei, fusiform and frontal eye field areas during simplified version of Sudoku puzzle (Qin et al., 2012). In fact, the same group of researchers had previously shown that the fMRI BOLD signal obtained from these regions of interest (ROI) can be used to predict the mental states during puzzle solving (Xiang et al., 2009). Their unique Sudoku paradigm (4×4 matrix, 2×2 subgrid)) eliminated the need of searching for the missing number or selecting the rules to be used. Rather, their paradigm was dedicated towards understanding the neural bases of basic heuristic problem solving processes in the knowledge domain. However, a complete Sudoku task includes both basic heuristic problem solving processes as well as searching & selecting the heuristic rules.

A 9 × 9 Sudoku requires the subject to be in sitting posture for some time, which limits the use traditional imaging techniques to study the brain activity during the task. In fMRI (functional magnetic resonance imaging) the subject has to be motionless and in a cage of magnets, this hardware restriction forms the ultimate limitation and restraints its’ use in claustrophobic participants. In addition, the supine motionless position is not the same used by an individual to solve the puzzle. The solution for such technical limitations is a perfectly non-invasive optical imaging technique called functional near infrared spectroscopy. Functional near infrared spectroscopy (fNIRS) is simple, safe and non-invasive neuroimaging technique of measuring brain activity (Ferrari and Quaresima, 2012). The principle behind the technique is to measure the absorbance of the infrared light to calculate the relative ratios of deoxygenated and oxygenated hemoglobin by modified Beer-Lambert law (Bunce et al., 2006). This hemodynamic response provides the indirect information about the brain activity as neural activation and vascular response are tightly coupled together, known as neurovascular coupling (Villringer and Chance, 1997). Another advantage of fNIRS over other imaging studies is its portability and robustness that makes it easy to study tasks akin to daily routine activities (Malik et al., 2017). The fNIRS fits the best technique to study puzzles like Sudoku. fNIRS system when applied to the hair free region on the forehead provides us with the activity of the PFC (Pre Frontal Cortex). It forms an important research query regarding the contribution of the PFC while solving Sudoku puzzle.

## 2. Materials & methods

### 2.1 Participants

This study was approved by institute ethical committee and in accordance with the Code of Ethics of the World Medical Association (Declaration of Helsinki 2000). All the participants were well informed about the nature and purpose of the study, and written informed consent was obtained from each. The participants with any known history of mental illness or neurodegenerative disorders like dementia or any history of brain trauma or surgery were not included in the study. Thus, thirty-three apparently healthy individuals (nine females, 28.88 ± 2.50 years) with normal or corrected to normal vision and with previous exposure to Sudoku task participated for the study. Handedness was assessed by Edinburgh Handedness inventory (31 participants were right handed) (Oldfield, 1971).

### 2.2 fNIRS imaging

Participants were seated comfortably in an armchair throughout the task while their prefrontal cortical activity was monitored using continuous-wave fNIR imaging system (Model 1100 BIOPAC Systems Inc.). The forehead was first cleaned by spirited cotton before the placing the sensor pad. This pad consisted of 4 sources and 10 detectors, which are configured to generate 16 measurement locations on the forehead called optodes (Figure 1a). Fp1, Fp2, and Fz position were marked using international 10-20 electrode placement system. The centerline of fNIRS sensor pad was first aligned according to Fz site with the horizontal axis coinciding with symmetry of the eyes and later it was adjusted in vertical plane so that the lower optodes corresponded to Fp1 and Fp2 locations. Such a position registers the 16 optodes locations as depicted in figure 1b (Ayaz et al., 2006). The fNIRS sensor pad was stabilized with an elastic band to prevent displacement. The source in the sensor pad has LED lights with peak wavelengths at 730nm and 850nm i.e. one above and other below the isosbestic point (∼800nm) where absorption spectrums of oxygenated and deoxygenated hemoglobin cross. The contact of each source and detector was ensured by observing the record online. Gain and power of LED current were accordingly adjusted to obtain optimal intensities for all the wavelengths. After stabilization of the record, 10-second pre-scan baseline was recorded before start of the task. Online data acquisition and visualization were done using COBI Studio software.

**Figure 1:**
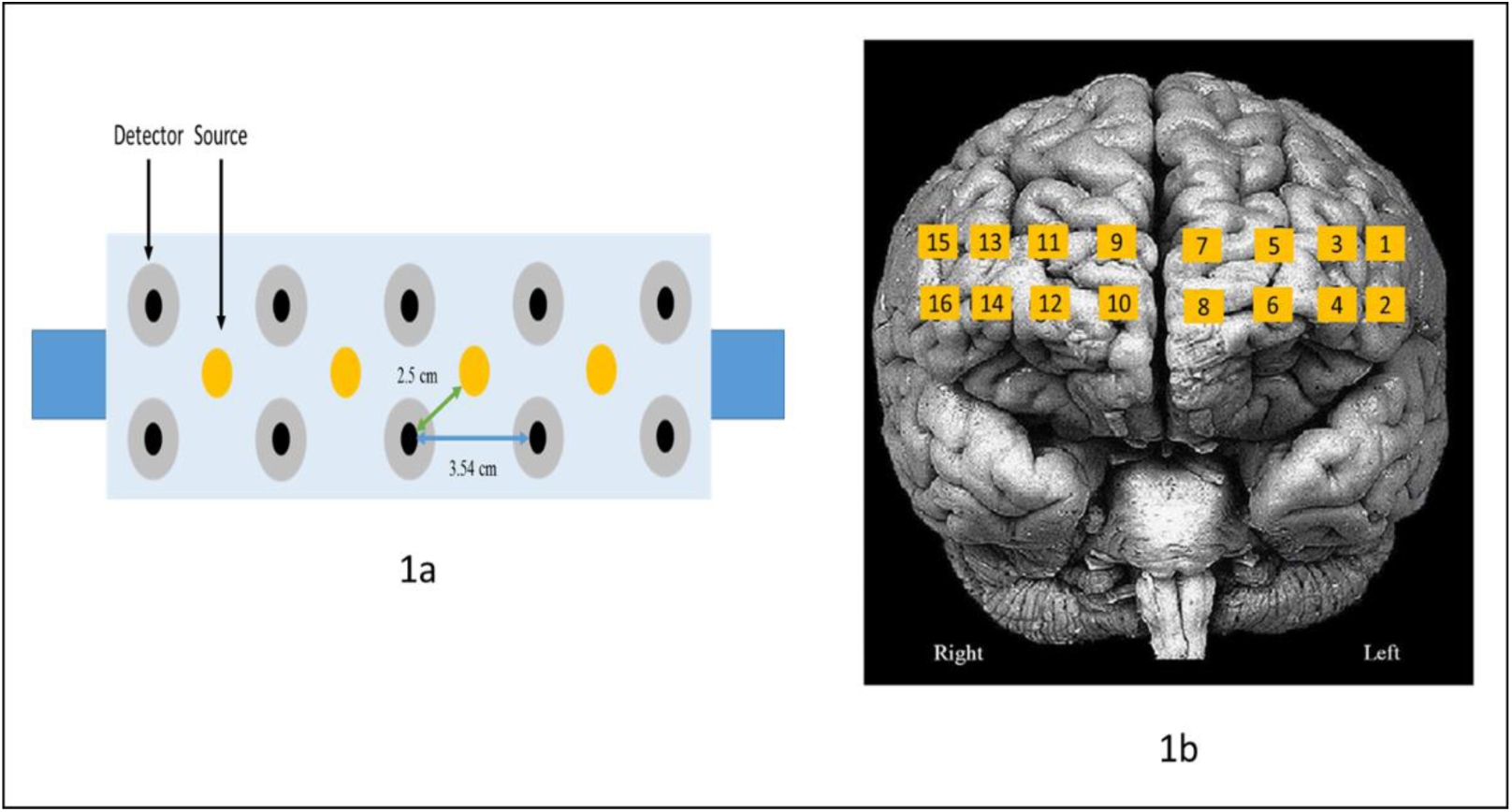
a. fNIRS Sensor Pad: - the sensor pad was placed on the forehead of the participants and aligned using 10-20 electrode placement system as mentioned in the text. The sensor pad consists of 4 sources and 10 detectors, which are configured to generate 16 optodes. b. Optode locations: 16 optode locations are depicted on brain surface frontal view.

### 2.3 Study Protocol

After familiarization with the lab setup and practice sessions, participants were asked to perform the task using paper-pen technique (akin to daily routine) with continuous recording of the prefrontal activity. Sudoku task was divided into two steps figure 2. During step 1, the participant had to fill one, two, or three missing numbers (1 to 9) in the 3 × 3 grids starting from the first row i.e. only subgrid rule to be used. The participants had to complete eighteen such grids (presented on the same paper sheet) with 33 total missing numbers for a period of 50 seconds. Step 2 consisted of an easy level Sudoku (9 × 9 matrix with subgrids of 3 × 3) composed from the 3 × 3 grids used in step 1. During step 2 apart from filling in the missing numbers the participant also has to keep in mind the row and column rule of the Sudoku puzzle. After stabilization of the fNIRS recording the task began with rest 1/baseline (40 seconds) → step 1 (50 seconds) → rest 2 (40 seconds) →step 2 (50 seconds) →rest 3 (40 seconds). These timings were decided from the results of the pilot study. Digital timer beeped at the end of each breaks and participant was instructed to change the sheet. Blank sheets were used during the rest periods.

**Figure 2:**
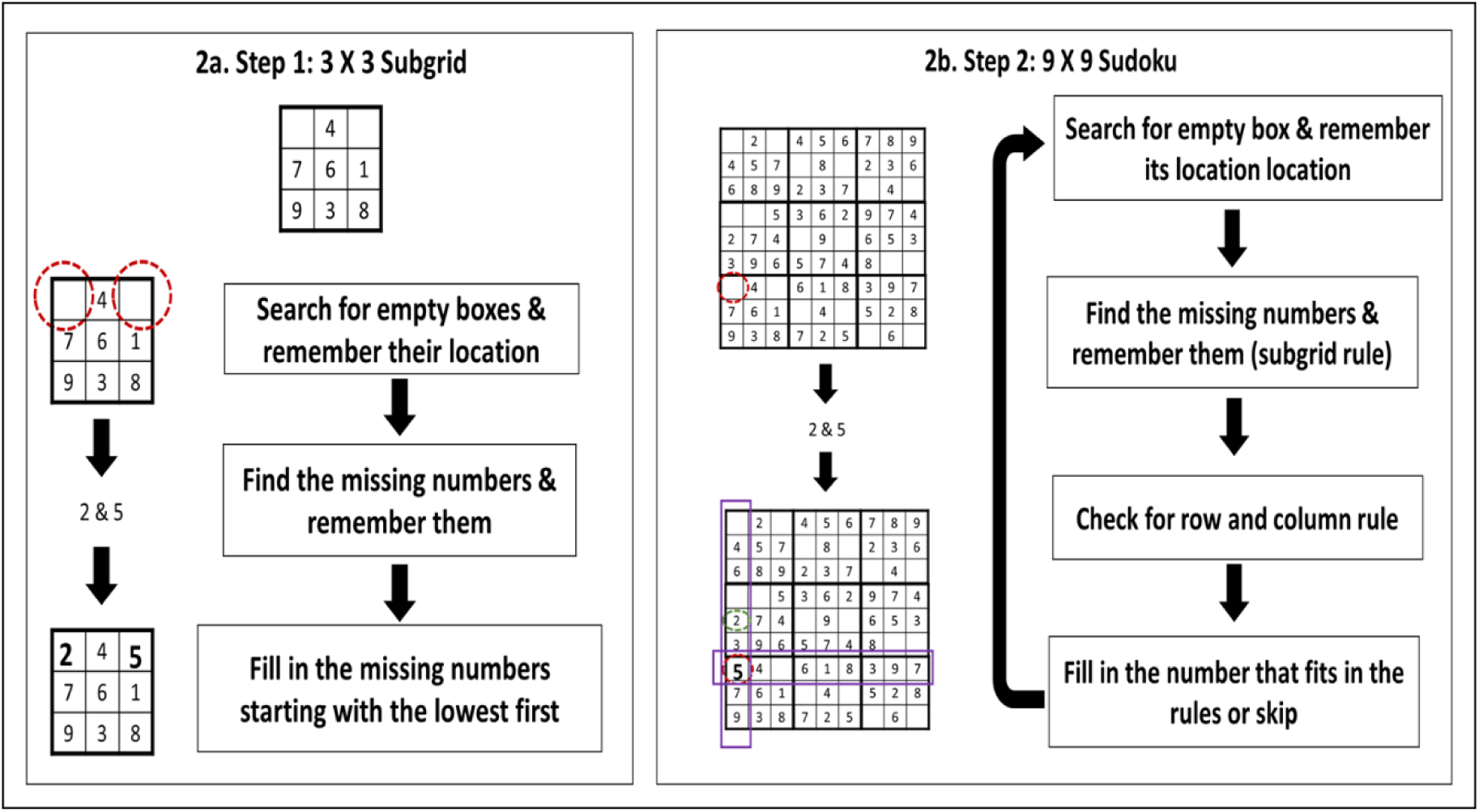
a. Step 1 i.e. 3×3 subgrid:- in this particular example the participants needs to fill in the missing numbers (2 & 5); b. Step 2 i.e. 9×9 subgrid:- here additionally the participants needs to fill in missing number according to row and column rule.

### 2.4 Data Analysis and Statistics

After data acquisition, analysis was done offline using fNIRsoft software. Concentrations of oxyhemoglobin and deoxy-hemoglobin were calculated using modified Beer-Lambert law with respected to prescan baseline of 10s. Various statistical methods have been developed to extract the neuronal related activity from the noisy signal. In this paper, we have used classical two-way ANOVA as well as General Linear Model based approach to analyze the data.

#### 2.4.1 Two-way ANOVA

The data was first subjected to low pass filter (0.40 Hz) with an order of 20 to remove trends due to respiratory, cardiovascular, or other high frequency artifacts. Two way ANOVA (3 task factors x 16 optodes) was used to compare the average concentration of oxyhemoglobin (HbO) and deoxy-hemoglobin (HbR) across the 16 optodes during baseline, step 1, and step 2. Post hoc analysis was done using Tukey’s multiple comparison test with alpha as 0.05.

#### 2.4.2 General Linear Model

Statistical analysis was also done using mass-univariate approach based on the General Linear Model in NIRS-SPM (Ye et al., 2009). The task design was used to create a square wave with rest periods having value of 0 and task periods as 1, which was then convoluted with hemodynamic response wave function to generate theoretical response. This theoretical response was compared with actual response during statistical analysis. Wavelet minimum description length detrending algorithm was applied to the data in order to remove trends due to respiratory, cardiovascular, or other experimental errors. This method prevents over/under fitting and facilitates optimal model order selection for the global trend estimate (Jang et al., 2009). Precoloring method with low pass hrf filter was used for corrections in temporal autocorrelation (Ye et al., 2009). Three contrast model matrices were assessed viz. contrast 1 (step 1-rest), contrast2 (step 2-rest), and contrast3 (step 2-step 1). The general linear model approach compared the covariance of each of the time points of the theoretical response to the actual response to create a t-test statistics for each channel.

## 3. Results

On visual inspection, data from three participants was excluded because either the optodes had saturated signal or there were motion artifacts shifting the baseline. Thus, we were able to record prefrontal activity in 28 right-handed participants (8 females, 23.04 ± 2.60 years). Results of the data obtained from 2 left-handed individuals are mentioned separately in supplementary data. The participants were able to solve step 1 and step 2 with an accuracy of 99.29 ± 1.50 % and 98.21 ± 2.64 % respectively. Figure 3a shows the grand average HbO and HbR levels across all optodes in all the participants. There was an increase in HbO levels (and decrease in HbR) during step 1 as well as during step 2. However, the HbO and HbR levels during the rest 2 period did not return to baseline values. Figure 3b & c shows the interpolated heat map of average HbO and HbR across all participants during the task. Furthermore, the step activity was divided into three parts viz. initial 10 seconds (where HbO and HbR levels were either increasing or decreasing), middle 30 seconds (where HbO and HbR levels were relatively stable), and late 10 seconds (where HbO and HbR levels were either decreasing or increasing). Similarly, the rest period was divided into two parts viz. initial 20 seconds and later 20 seconds.

**Figure 3:**
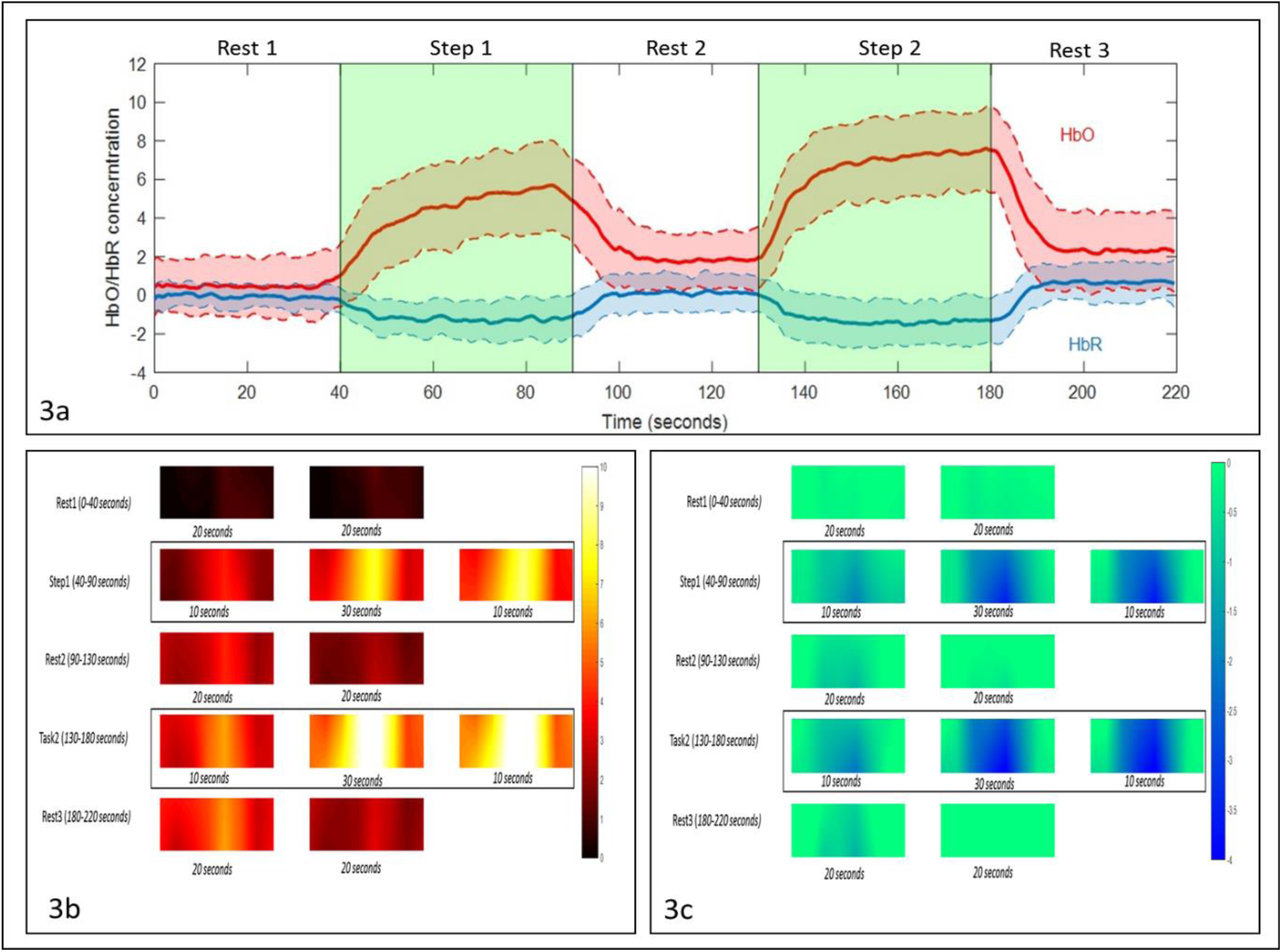
a. The solid red and blue lines represent the grand average HbO & HbR concentrations across all optodes during the task respectively. The shaded red & blue regions represent the standard deviation of HbO & HbR concentrations during the task respectively. b. & c. Interpolated heat map of average HbO and HbR across all participants during the task

We calculated the relative changes in HbO and HbR concentrations during step 1 and step 2 by subtracting the pre-step rest activity. Average concentrations of HbO and HbR during step 1 and step 2 were calculated using a middle stable portion. Figure 4a &b depicts grand average concentrations of HbO & HbR levels during baseline, step 1, and step 2 at 16 optodes locations for all participants.

**Figure 4:**
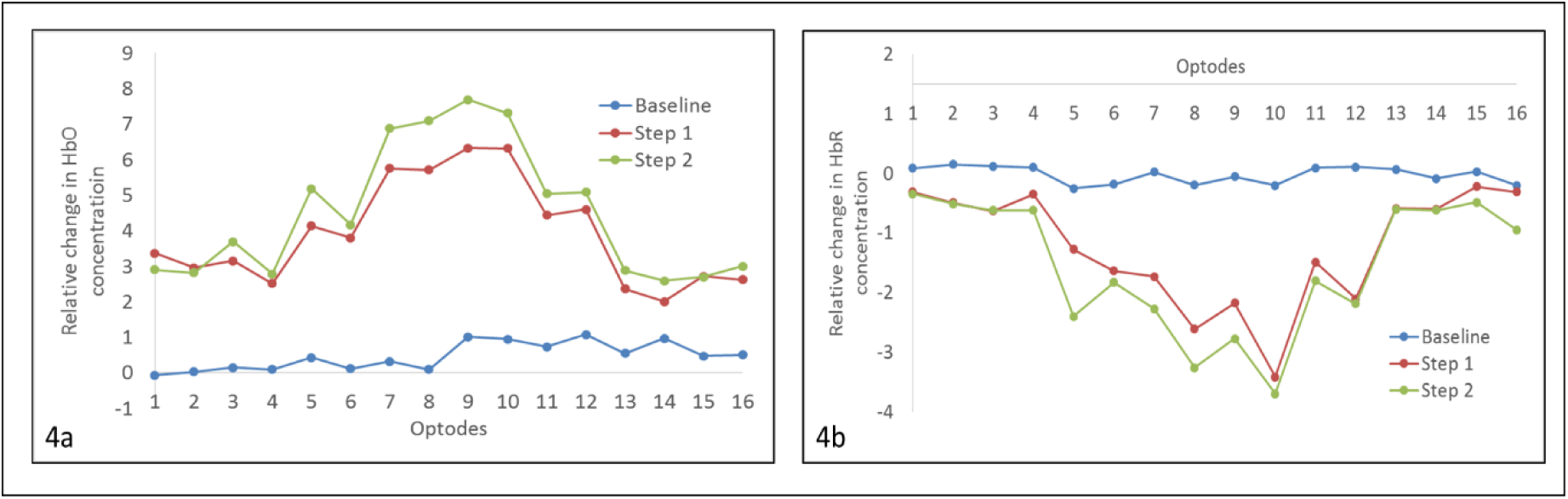
Grand average concentrations of HbO & HbR levels during baseline, step 1, and step 2 at 16 optodes locations for all participants

Two-way ANOVA was conducted that examined the effect of the task (baseline, step 1 and step 2) and 16 optodes locations on HbO and HbR. There was a statistically significant interaction between task and optode locations on HbO, F (30, 1296) = 19.03, p<0.0001 and HbR, F (30, 1296) = 41.91, p<0.0001. Post hoc analysis showed a significant increase in relative HbO concentrations at all optodes during step 1 and step 2 as compared to baseline (p<0.0001). In addition, there was a significant increase in relative HbO concentrations at optodes 5, 7, 8, 9, 10 during step 2 as compared to step 1 (Table 1). HbR levels were significantly decreased at all optodes during step 1 and step 2 as compared to baseline (p<0.0001). Moreover, HbR levels were significantly decreased at optode 4, 5, 7, 8, 9, 10, 11, 15, and 16 during step 2 as compared to step 1(Table 1).

**Table 1:** Adjusted p value at various optode locations after post hoc analysis (Step1 vs. Step2 for HbO and HbR)

|  |  | Optode 1 | Optode 2 | Optode 3 | Optode 4 | Optode 5 | Optode 6 | Optode 7 | Optode 8 |
| --- | --- | --- | --- | --- | --- | --- | --- | --- | --- |
| Step 1 vs. Step 2 | HbO | 0.2786 | 0.8849 | 0.1899 | 0.6629 | **0.0019** | 0.4658 | **0.0007** | **<0.0001** |
|  | HbR | 0.6733 | 0.8703 | 0.9228 | **<0.0001** | **<0.0001** | **<0.0001** | **<0.0001** | **<0.0001** |
|  |  | <u>Optode 9</u> | <u>Optode 10</u> | <u>Optode 11</u> | <u>Optode 12</u> | <u>Optode 13</u> | <u>Optode 14</u> | <u>Optode 15</u> | <u>Optode 16</u> |
| <u>Step 1 vs. Step 2</u> | <u>HbO</u> | **<0.0001** | **0.0037** | 0.1219 | 0.2438 | 0.2074 | 0.135 | 0.9975 | 0.4406 |
|  | <u>HbR</u> | **<0.0001** | **<0.0001** | **<0.0001** | 0.0958 | 0.9635 | 0.8144 | **<0.0001** | **<0.0001** |

Group analysis done using General Linear Model in NIRS-SPM depicts the statistical significant t maps for HbO and HbR for three contrast models viz. contrast 1 (step 1 - rest), contrast 2 (step 2 - rest), and contrast 3 (step 2 - step 1) (Figure 5). There was significant activation seen at all optode locations during step 1 and step 2 as compared to the rest (contrast 1 and contrast 2). In addition, there was a significant activation seen during step 2 as compared to step 1 (contrast 3) in medial PFC regions corresponding to optodes 7, 8, 9, 10.

**Figure 5:**
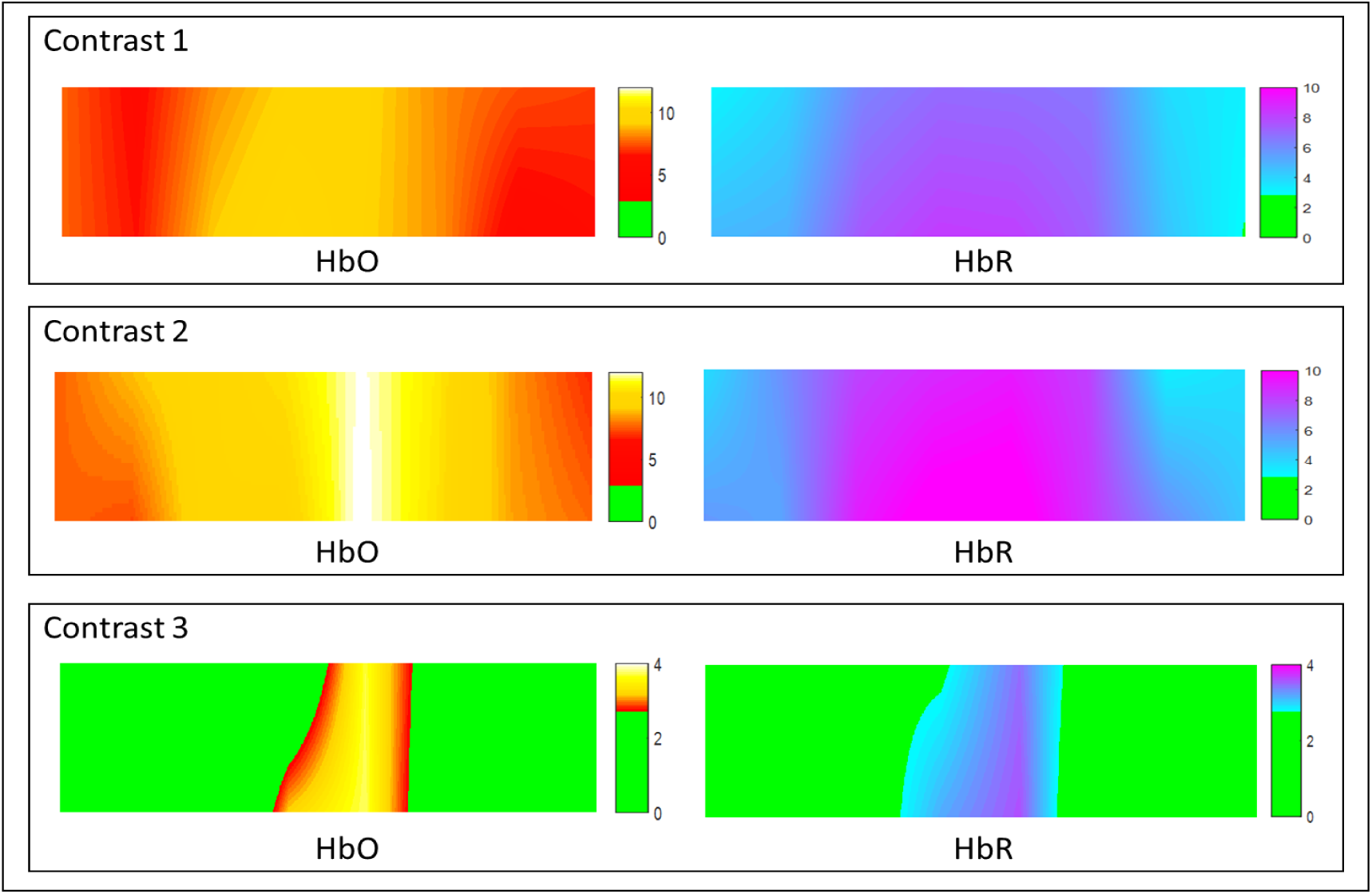
Statistically significant interpolated t maps for HbO and HbR for three contrast models viz. contrast 1 (step 1 - rest), contrast 2 (step 2 - rest), and contrast 3 (step 2 - step 1). p-value: 0.05 & its corresponding z-threshold: 2.69.

## 4. Discussion

In this study, we were able to record the prefrontal cortical activity during Sudoku task in 28 right-handed participants and two left handed participants. To our best knowledge, this is first fNIRS study that tried to unravel the role of PFC during Sudoku task. In order to achieve this objective, the Sudoku task was uniquely divided into two steps that tried to separate the complex heuristic rule selection processes. Step 1 in our task eliminated the need for selecting a row or column rule. Thus, PFC activity during step 1 represents simple processes like searching empty boxes, remembering their location, finding the missing numbers, remembering the numbers, and placing the numbers while ignoring the stimulus coming from nearby subgrids. Using the same grids to create a simple level Sudoku (Step 2) will now involve additional processes like searching and applying the column & row rules. That is the step 2 involves more complex heuristic rule selection processes. Thus, the increased PFC activity during Step 2 represents these additional processes. As PFC is involved in many cognitive functions like working memory, attention and decision making (Bench et al., 1993; Euston et al., 2012; Kane and Engle, 2002; Rueckert et al., 1996). Thus, it is not surprising to find increased PFC activity during both steps of Sudoku task. Step 1 in our study is can be considered similar to the simple heuristic task and step 2 is similar to the complex heuristic task used by Wang et al. These researchers have shown that such complex tasks cost more time than simple tasks as cognitive processes involved in heuristic processing is complex (Wang et al., 2009). There is significant activation of left PFC during the complex task as compare to simple tasks (Long et al., 2012). In this study, we saw that step2-step1 contrast had medial regions activated. Indeed, these medial regions correspond to part of frontopolar cortex (FPC) which is proposed to be higher within the PFC hierarchy (Boschin et al., 2015). FPC is involved in behaviors that include complex decision making (Daw et al., 2006), prospective memory (Okuda et al., 2007) and rule-specific cognitive processing (Sakai and Passingham, 2006). In addition, activity in medial prefrontal regions decreases if trials become familiar (Zhou et al., 2012). Thus, increased activity in medial regions during step 2 minus step 1 represents the processes involved in Sudoku task and not due to familiarity. In other sense, it can be said that Sudoku as a leisure time cognitive stimulation activity helps in differentially activating the PFC regions.

Data acquisition and analysis methods used for understanding task related fNIRS signal are in their adolescent period as compared to fMRI analysis methods. In this study, we used both classical ANOVA and GLM methods to analyze the data in order to give convergent evidence of PFC activity during paper-pen block Sudoku task. However, the GLM approach supersedes the classical technique in reducing false negative and false positive results (Tachtsidis and Scholkmann, 2016). Nevertheless, newer fNIRS devices need to be used to rule out effect of extra-cerebral hemodynamic changes, which can be achieved by adding short-channel regression techniques (Goodwin et al., 2014). Moreover, future studies can be directed towards recording of the complete cortex using multi-optode fNIRS. Such recording will provide us with wider view of the networks involved during complex 9×9 Sudoku task processing. Future studies may also be directed towards study of performance during Sudoku task. Such explorations will help us to device newer tasks in elderly or in patients requiring neuro-rehabilitation.

To conclude, there is significant involvement of PFC during Sudoku task. Both the medial and lateral regions of PFC are activated when the participant is solving the Sudoku task. However, the medial regions of PFC play a differential role, especially when we consider the row and the column rule of Sudoku.

## Acknowledgements

No other financial disclosures or potential conflicts of interest related to this article are reported.

## Supplementary Data

We could also record the PFC activity during Sudoku task in two left-handed participants (age 21 and 25 years, both males). There completed both step 1 and step 2 with accuracy of 100 %. Figure 6 depicts the grand average HbO and HbR levels across all optodes in the left-handed participants. Similar to the result obtained in other participants there was an increase in HbO levels (and decrease in HbR) during step 1 as well as during step 2. However, the HbO and HbR levels during the rest 2 period did not return to baseline values. Figure 6b & c shows the interpolated heat map of average HbO and HbR across left-handed participants during the task. Furthermore, the step activity and the rest period were divided into 3 and 2 parts respectively similar to the record obtained in other participants.

**Supplementary Figure 6:**
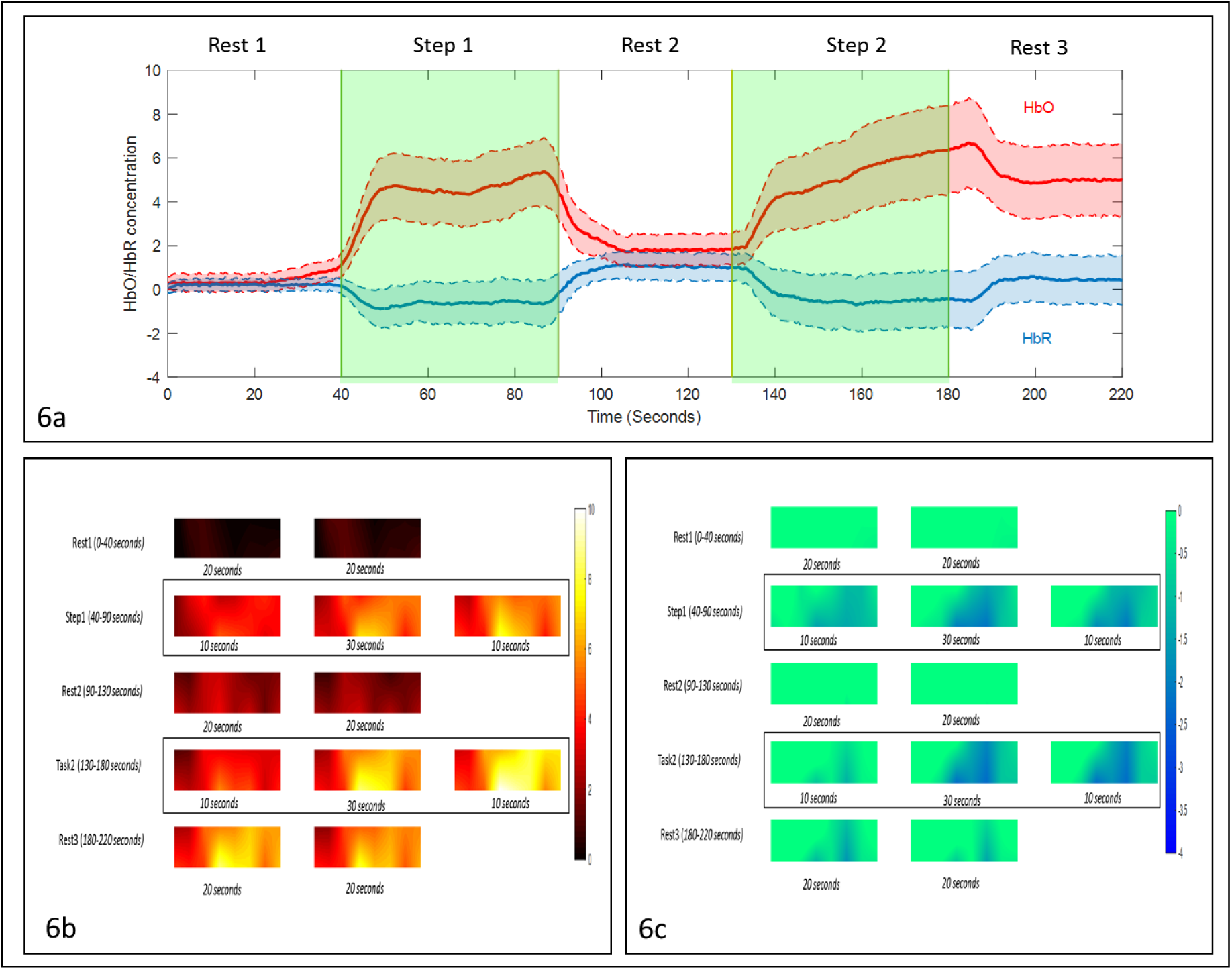
a. The solid red and blue lines represent the grand average HbO & HbR concentrations across all optodes during the task respectively. The shaded red & blue regions represent the standard deviation of HbO & HbR concentrations during the task respectively. b. & c. Interpolated heat map of average HbO and HbR across all participants during the task

